# Optimizing guide RNA selection and CRISPR/Cas9 methodology for efficient generation of deletions in *C. elegans*

**DOI:** 10.1101/359588

**Authors:** Vinci Au, Erica Li-Leger, Greta Raymant, Stephane Flibotte, George Chen, Kiana Martin, Lisa Fernando, Claudia Doell, Federico I. Rosell, Su Wang, Mark L. Edgley, Ann E. Rougvie, Harald Hutter, Donald G. Moerman

## Abstract

The *Caenorhabditis elegans Gene* Knockout (KO) Consortium is tasked with obtaining null mutations in each of the more than 20,000 open reading frames (ORFs) of this organism. To date, approximately15,000 ORFs have associated putative null alleles. A directed approach using CRISPR/Cas9 methodology is the most promising technique to complete the task. While there has been substantial success in using CRISPR/Cas9 in *C. elegans*, there has been little emphasis on optimizing the method for generating large insertions/deletions in this organism. To enhance the efficiency of using CRISPR/Cas9 to generate gene knockouts in *C. elegans* we have developed an online species-specific guide RNA selection tool (http://genome.sfu.ca/crispr). When coupled with previously developed selection vectors, optimization for homology arm length, and the use of purified Cas9 protein, we demonstrate a robust, efficient and effective protocol for generating deletions. Debate and speculation in the larger scientific community about off-target effects due to non-specific Cas9 cutting has prompted us to investigate through whole genome sequencing the occurrence of single nucleotide variants and indels accompanying targeted deletions. We did not detect any off-site variants above the natural spontaneous mutation rate and therefore conclude this modified protocol does not generate off-target events to any significant degree in *C. elegans.*

## Introduction

CRISPR/Cas9 is the current technology of choice for genome editing (Sander and Joung, 2014). This is due to its versatility, as this is an RNA-guided system where a 20-base guide RNA (crRNA) directs a Cas9 nuclease (from *Streptococcus pyogenes*) to the target sequence providing high specificity and minimal off-target site effects. The endonuclease makes a double-strand cut at the target site (Gasiunas *et al*. 2012; Jinek *et al*. 2012; Jiang *et al*. 2013), which can then be repaired through Non-Homologous End Joining (NHEJ) or Homology-Directed Repair (HDR). CRISPR/Cas9 technology was first adapted for *C. elegans* in 2013 (Cho *et al*. 2013; Friedland *et al*. 2013; Chiu *et al*. 2013; Chen *et al*. 2013; Dickinson *et al*. 2013; Tzur *et al*. 2013; Lo *et al*. 2013; Katic *et al*. 2013; Waaijers *et al*. 2013) and since then the community has produced increasingly sophisticated methods to mutate, delete and tag genes (for example, Dickinson *et al*. 2015; Norris *et al*. 2015; Paix *et al*. 2014, 2016).

Our facility, along with several international laboratories, is attempting to isolate deletion alleles for the majority of genes in *C. elegans*. We have tested a number of the current methodologies for the CRISPR/Cas9 system and have successfully isolated several small deletions. We found that none of the CRISPR/Cas9 protocols presently available to the worm community were suitable, as is, for an ambitious high-throughput approach, which is what the *C. elegans* Gene Knockout (KO) Facility requires. In this paper we examine the details of several parts of the method with the aim of obtaining a more efficient experimental procedure that may in turn lead to a greater yield of deletions in a timely manner. These parts are (1) the repair mechanism and integrant selection; (2) homology arm length; (3) Cas9 delivery; and (4) guide RNA selection.

We first examined repair mechanisms and selection procedures. While one can obtain deletions utilizing NHEJ, which is reportedly polymerase theta-mediated repair in the *C. elegans* germline (van Schendel *et al*. 2015), the most versatile methodologies use HDR to introduce designer modifications at precise locations in the genome (Yang *et al*. 2013; Auer *et al*. 2014; Port *et al*. 2014; Bottcher *et al*. 2014; DiCarlo *et al*. 2013; Kim *et al*. 2014; Arribere *et al*. 2014; Zhao *et al*. 2014). HDR is homologous recombination generated at the Cas9 cut site that is facilitated by exogenous DNA with homology to the regions flanking the cut site in the genome. Two recent papers coupled HDR in *C. elegans* with the introduction of drug selection for either hygromycin (Dickinson *et al*. 2015) or G418 (Norris *et al*. 2015) as part of the screening process. This is an important modification as drug selection improves throughput by greatly reducing the number of animals that need to be screened. For the studies described in this paper, we have opted to use the G418 protocol described by Norris *et al*. (2015). Their protocol requires the introduction of a repair template containing a dual-selection cassette: a G418 resistance gene (*neoR*) and a pharyngeal GFP reporter (P*myo-2::GFP*), flanked by long homology arms (Figure 1). As described earlier, the drug resistance marker allows F1 selection on G418 and reduces primary screening efforts by eliminating animals that do not carry the repair template. Integration of the repair template via HDR allows the simultaneous deletion of a defined genomic region and introduction of the dual-selection cassette. Furthermore, the GFP marker makes it possible to discern recombinant animals from those harboring concatemer arrays. Recombinant animals display dim and uniform expression of GFP in the pharyngeal muscle, whereas concatemer array expression is often bright and mosaic. Additional pharyngeal and body-wall RFP-bearing plasmids (P*myo-2::RFP* and P*myo-3::RFP*, respectively) aid in the identification of non-integrants since these independent plasmids should only be expressed from concatemer arrays. Another convenient feature of this vector is the ability to easily excise the selection cassette using Cre recombinase as there are *LoxP* sites flanking the cassette (Figure 1; Norris *et al*. 2015).

**Figure 1.**
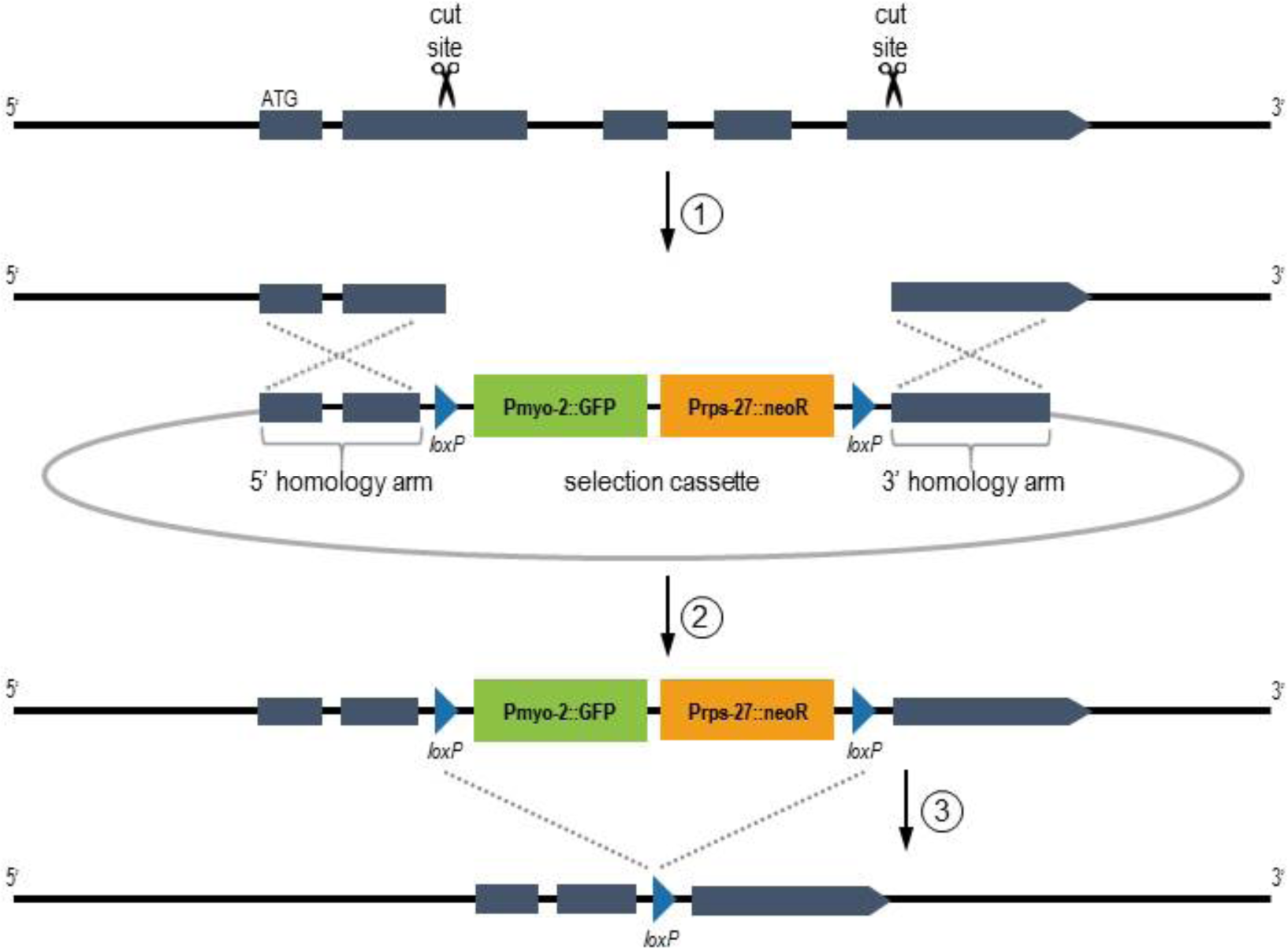
Generation of a deletion using the CRISPR/Cas9 protocol. (1) Guide RNAs direct Cas9 to create targeted DSBs in the gene of interest. (2) Through HDR, a portion of the ORF is replaced with the selection cassette containing pharyngeal GFP and G418 resistance markers. (3) The dual-marker cassette, flanked by *loxP* sites, can be excised from the genome by injecting Cre recombinase. (Adapted from Norris *et al*. 2015).

Once we settled on an HDR procedure, accompanied by drug selection, we then examined the length of homology arm required for efficient HDR. The optimal length of homology arm required for efficient modification via HDR at the target site is still a matter for debate. A recent paper by Paix *et al*. (2014) has shown that short, 30 to 60 nucleotide, homologous linear repair templates can be highly efficient for gene editing over small target intervals. In contrast, two papers (Dickinson *et al*. 2015; Norris *et al*. 2015) have shown that large inserts (>1.5 kb) require longer homology arms (>500bp).

We next examined how best to deliver Cas9 to facilitate efficient DNA cutting. There are at least three ways to deliver Cas9 to the target DNA; via plasmid, as messenger RNA or as purified protein. The Seydoux lab has shown that the third method, direct injection of Cas9 protein as a ribonucleoprotein (RNP) complex (Cas9 protein, the structural tracrRNA and the target-specific crRNA), is fivefold more efficient than injection of plasmids coding for these reagents in *C. elegans* (Paix *et al*. 2014). This advance could decrease injection time considerably and eliminate the need to generate guide RNA expression plasmids for every gene target since they could be synthesized.

We recognized that the selection of effective guide RNAs in *C. elegans* to efficiently obtain deletion mutations is critical. Although there is no clear consensus for effective guide RNA design constraints, certain metrics for efficacy have been employed by various researchers (reviewed in Mohr *et al*. 2016). We reasoned that our overall CRISPR/Cas9 success rate could be substantially improved by the application of a set of standard filters to the entire complement of all possible guide RNAs in the *C. elegans* genome, as a type of pre-selection to eliminate from consideration guide RNAs that could be expected to perform poorly. These filters are based on accumulated observations from many organisms. Our online species-specific guide RNA selection too is available at (http://genome.sfu.ca/crispr).

In this manuscript we present experimental data for each of these parts of the method, leading to a highly efficient standard protocol for using CRISPR/Cas9 to generate deletions in *C. elegans*. We describe our new tool for the design and selection of guide RNAs specific to *C. elegans*; test the length of homology arms necessary and sufficient for using CRISPR/Cas9 to engineer deletions of several hundred nucleotides while introducing a large (5.4 kb) selection cassette; and compare the relative on-site efficiency of delivering Cas9 encoded in a plasmid to the direct delivery of Cas9 protein using a number of different gene targets. As our facility provides deletion strains to the Caenorhabditis Genetics Center (CGC), which then supplies the larger worm community, we want to ensure, if possible, that there are no additional mutations in these CRISPR/Cas9-generated events. To this end, we performed WGS (whole genome sequencing) on several of our deletion strains to determine the frequency of off-site mutations and to examine rearrangements at the target gene itself.

## Materials and methods

**Designing a *C. elegans-*specific CRISPR guide RNA selection tool:** Guide RNAs have been designed for the whole *C. elegans* genome and made available to the research community (see data availability section). Briefly, a series of filters on the potential guide sequences have been applied with an in-house Perl script using the reference genome sequence and gene information from WormBase version WS250 (http://www.wormbase.org). In a first step, for each Protospacer Adjacent Motif (PAM) site in the genome we kept the corresponding adjacent 20-base guide only if its GC content was between 20% and 80% and no poly-T tracts of length 5 or longer were present. We annotated the presence or absence of the sequence GG at the 3’ end of each guide since guides ending with GG are expected to have higher efficiency for 3’ NGG PAMs (Farboud and Meyer, 2015). Guides for which the seed region, defined as 12 bases at the 3’ end plus the PAM, was not unique in the genome were eliminated. The uniqueness of those 15-mers was assessed with an in-house C code modified from Flibotte and Moerman (2008). The guide plus PAM sequences were then aligned to the whole genome with bwa aln (Li and Durbin, 2009) allowing an edit distance of 3 and guides mapping to multiple locations in the genome were eliminated. The minimum free energy in kcal/mol was calculated for each guide with the program hybrid-ss-min (Markham, 2003) to help the researchers assess potential self-folding of the RNA. The location of the cut site associated with each guide was then annotated according to the gene feature being targeted, if any. All the guides have been loaded into a database, which is searchable via a web interface. Users can search the database by entering a genomic interval or by searching a gene name if they wish to find guides that have cut sites within genes. Constraints can be applied to the GC content, folding energy, and the presence of GG at the 3’ end of the guides being returned.

**CRISPR/Cas9 deletion procedures:** CRISPR/Cas9 edits were generated using a protocol previously described by Norris *et al*. (2015), with some modifications. Repair templates were assembled using the NEBuilder Hifi DNA Assembly Kit (New England BioLabs) to incorporate homology arms into a dual-marker selection cassette (*loxP* + P*myo-2::GFP::unc-54* 3’UTR + P*rps-27::neoR::unc-54* 3’UTR *+ loxP* vector provided by John Calarco). Homology arms were ordered as DNA gBlocks from Integrated DNA Technologies (IDT), approximately 500 bp in length including 50 bp of overhang adaptor sequences for the assembly reaction.

Depending on the experiment, CRISPR/Cas9 edits were generated using one or two guide RNAs, and Cas9 was delivered via one of two methods: a Cas9 expression plasmid (Fu *et al*. 2014) or purified Cas9 protein (Paix *et al*. 2015). For experiments using the Cas9-expression plasmid, a single guide RNA (sgRNA) vector was constructed by performing PCR on a p*U6::klp-12* vector (Friedland *et al*. 2013), to incorporate the 20-bp guide RNA sequence via the forward primer. Injection mixes consisted of: 100 ng/µL sgRNA plasmid, 50 ng/µL repair template, 50 ng/µL Cas9 expression plasmid (P*eft- 3::Cas9::NLS SV40::NLS::tbb-2 3’UTR)*, 5 ng/µL pCFJ104 (P*myo-3::mCherry*), and 2.5 ng/µL pCFJ90 (P*myo-2::mCherry*). For Cas9 protein injections, crRNA and tracrRNA (synthesized by IDT) were duplexed and then combined into a Cas9 RNP complex following the manufacturer’s recommendations. Cas9 protein was purified according to the protocol described by Paix *et al*. (2015) from plasmid nm2973 (Fu *et al*. 2014) and stored at 34 µg/µL in 20 mM Hepes buffer pH 8.0, 500 mM KCl and 20% glycerol. Injection mixes consisted of 50 ng/µL repair template, 1.5 µM RNP complex, 5 ng/µL pCFJ104, and 2.5 ng/µL pCFJ90.

For each CRISPR target, approximately 32 young adult VC3504 (Moerman lab derivative of N2) hermaphrodites were injected as previously described by Kadandale *et al*. (2009) and screened according to the protocol in Norris *et al*. (2015).

**Whole-Genome sequencing (WGS) and analysis to measure off-target effects after CRISPR/Cas9 treatment:** Off-target effects were assessed using WGS for eight CRISPR/Cas9-derived strains generated for two genes from the same parental VC3504 population. Two guide RNAs were used for each of the target genes, *lgc-45* and *C34D4.2*. For the *lgc-45* gene we used guide sequences: TCCGATTCGAGTGGTTCACG and AAACCAATACGACCCATTCT. For the gene *C34D4.2* >we used guide sequences: CGCCTATCAAGTCCTCCACC and AGATTTTGGTTAGATTTCGA. To assess possible differences in off-target effects between Cas9 plasmid and Cas9 protein, we injected approximately 32 animals with either plasmid or protein for each set of CRISPR guides. All independently generated strains were sequence validated by PCR for correct insertion of the selection cassette. For each target, four independently generated strains (two from plasmid injections, and two from protein injections) were selected for WGS. We also sequenced the parental strain VC3504. All CRISPR strains were no more than three generations away from the parental strain when injected.

*C. elegans* strains were grown as described by Brenner (1974). Strains were allowed to grow on either 100 mm or 60 mm agar plates seeded with OP50 until just starved. Worms were washed off plates with sterile distilled water into 15 mL centrifuge tubes, pelleted, and washed with several changes of dH2O to remove residual *Escherichia coli* and other possible contaminants. The supernatant was removed after the final wash and a dense pellet of worms was transferred into a 1.5 mL freezer tube. The dense pellets of worms were frozen at -80 C and subsequently subjected to standard genomic DNA extraction. The extraction begins with Proteinase K digestion and treatment with RNAse A, followed by 6M NaCl protein precipitation. The DNA is then precipitated in isopropanol, washed in 70% ethanol, and finally resuspended in distilled water. The genomic DNA was run on 1% agarose gel and high molecular weight DNA was extracted using NEB’s Monarch Gel Extraction Kit. Sequencing libraries were made using Illumina’s Nextera XT library prep kit and run on an Agilent Bioanalyzer to check average fragment size. All strains were sequenced on MiSeq 2×300 or 2×75 runs. In one case, strain VC3823, the second read was unusable, so it was treated as a single run (i.e. not paired) at the analysis stage.

Sequence reads were mapped to the *C. elegans* reference genome version WS260 (http://www.wormbase.org) using the short-read aligner BWA version 0.7.16 (Li and Durbin, 2009). For each sample, this resulted in an average sequencing depth ranging from 22x to 41x with a median of 32x. Single-nucleotide variants (SNVs) and small insertions/deletions (indels) were identified and filtered with the help of the SAMtools toolbox version 1.6 (Li *et al*. 2009). Candidate variants at genomic locations for which the parental N2 strain VC3504 had a read depth lower than 10 reads or an agreement rate with the reference genome lower than 98% were eliminated from further consideration. Furthermore, for each of the eight CRISPR strains sequenced, only variants at locations where all of the other seven CRISPR strains agreed with the reference for at least 95% of the reads were kept. This last filter principally eliminated heterozygous candidates likely due to PCR artefacts (often in or adjacent to homopolymer stretches of A or T) or potentially present at low level in the parental population but undetected by our sequencing of VC3504. Each variant was annotated with a custom Perl script and gene information downloaded from WormBase version WS260. The read alignments in the regions of candidate variants were visually inspected with Integrative Genomics Viewer (IGV; Robinson *et al*. 2011; Thorvaldsdottir *et al*. 2013). Copy numbers were estimated from the alignments with a procedure analogous to that of Itani *et al*. (2015) using 1-kb wide sliding windows with the alignments from the parental strain as the reference in order to search for potential off-target genomic rearrangements.

**Quality Control testing of CRISPR/Cas9 target deletions using the Polymerase Chain Reaction (PCR):** WGS is an extremely informative method for determining the structure of a CRISPR mutation, but it is more labour-intensive, time-consuming and costly than PCR. We developed a straightforward PCR protocol as an efficient screening tool, which provides basic information about the selection cassette insertion and the desired deletion in four reactions per generated mutation. Two primer pairs are used to amplify the upstream and downstream insertion sites, each pair with one primer in the genomic sequence adjacent to the insertion and the other within the cassette sequence. Another pair of primers is used to amplify wild type (WT) sequence in the region of the deletion from both N2 and mutant templates. The latter primer pair is sometimes designed as a pair flanking the deletion, and sometimes as a pair with one primer flanking and one internal to the deleted region. There are four possible results from these PCRs; (A) The WT product is present and of the correct size from N2 and absent from the mutant, and both insertion-site products are present and of the correct size; (B’ and B”) The WT assays are correct and one insertion-site product is correct but the other one is missing or of incorrect size; (C) The WT assays are correct and both insertion-site products are absent or of incorrect size; and (D) The WT assay on the mutant is incorrect. For cases A-C, we conclude that the gene is disrupted and that the insertion sites are correct or rearranged. For case D, where a WT product of the size predicted for N2 is produced from the mutant template, we conclude that the gene is not disrupted.

**Data Availability:** The raw sequence data from this study have been submitted to the NCBI BioProject (http://www.ncbi.nlm.nih.gov/bioproject) under accession number PRJNA 473363 and can be accessed from the Sequence Read Archive (SRA; https://www.ncbi.nlm.nih.gov/sra) with accession number SRP149097. CRISPR guide RNA sequences designed for the whole genome are available using the search tool at http://genome.sfu.ca/crispr/ and can be viewed interactively within JBrowse at WormBase (http://www.wormbase.org) by activating the track called “CRISPR_Cas9 sgRNA predictions”. All the guides can also be downloaded in bulk as a bed file at http://genome.sfu.ca/crispr/

## Results

### A website for guide RNA design and selection

In the context of a production laboratory like our gene knockout facility it is crucial to use the time of the laboratory personnel effectively and eliminate unnecessary tasks. With that in mind, we attempted to streamline the design of guide RNAs by pre-calculating, pre-filtering and annotating potential guide RNAs across the whole *C. elegans* genome (see Materials and Methods for details). An important feature of the guide RNA design site is that it includes full gene annotation such that guide RNAs are identified in relation to the gene, which allows judicious choices to be made depending on the goal of the experimental procedure. Since such a resource is potentially useful to the research community, we decided to make our database of guide RNAs searchable via a public web site (http://genome.sfu.ca/crispr; Figure 2A). The same set of guides are also viewable as a JBrowse track, CRISPR/Cas9 sgRNA predictions, at WormBase (http://www.wormbase.org; Figure 2B), or within IGV (Robinson *et al*. 2011; Thorvaldsdottir *et al*. 2013) after downloading a bed file available on our guide RNA web site.

**Figure 2.**
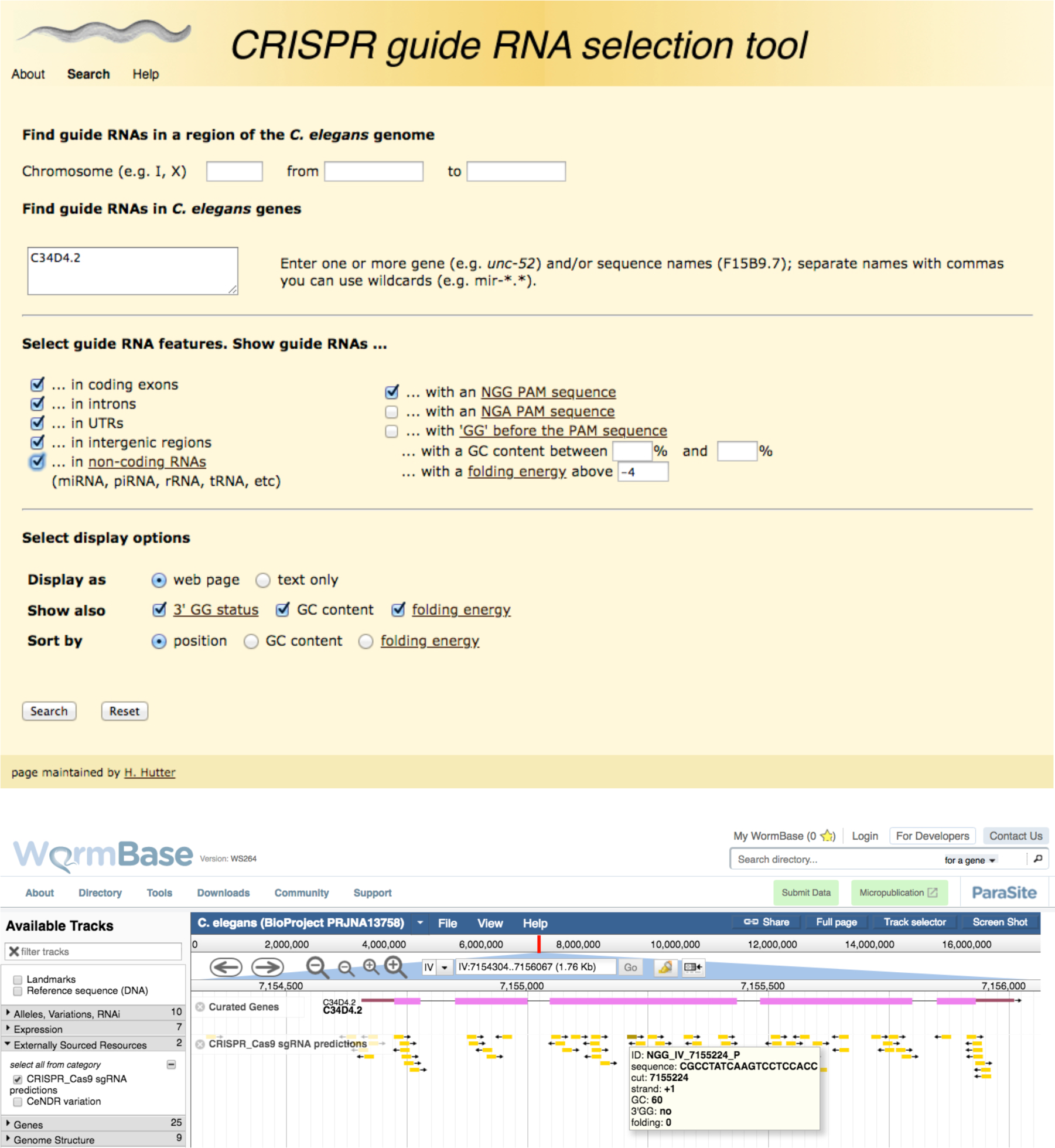
A and B. The CRISPR guide RNA selection tool. In panel A is a screenshot of the website for designing guide RNAs. Panel B illustrates guide RNA predictions from the design site available at WormBase and shows them along the structure of a gene.

As expected, the use of the guide RNA selection tool significantly reduced the amount of time spent at the design stage by laboratory personnel, but most importantly, and somewhat unexpectedly, this improved the overall laboratory throughput by promoting the selection of more efficient guides. Our current success rate for obtaining deletions using these guides is approximately 85% (based on 112 unique deletion attempts), with the vast majority of successes achieved during a first round of 32 injections (divided 4 worms per plate). Filtering and selecting for unique guide RNAs is most likely the major contributor to the negligible off-target background we observe after CRISPR/Cas9 use (see below).

### Homology–directed repair (HDR) and homology arm length

HDR allows DNA fragments to be incorporated into a precise region of the *C. elegans* genome. Homology arms flanking these DNA fragments dictate the genomic region that will be simultaneously deleted (if so desired) and replaced. These homologous regions also must be immediately adjacent to the Cas9 double-strand cut site in the genome. To date, the homology arm length used in different protocols has varied from 35bp (Paix *et al*. 2014) up to approximately 2 kb (Dickinson *et al*. 2015; Norris *et al*. 2015). Paix *et al*. (2016) report optimal HDR efficiency with 35-bp homology arms on a single-stranded template for making small edits. We conducted a series of experiments to test the efficiency of generating a 536 bp deletion and inserting a large selection cassette (5.4 kb) at a Cas9-induced double-strand break using a single guide (5’- AGGAGTTGGGAAAAGCGCAT-3’) with different lengths of homology arms (Table 1). The length of upstream and downstream homology arms differed slightly due to synthesis constraints on the sequence of gBlocks ordered from IDT.

**Table 1.**
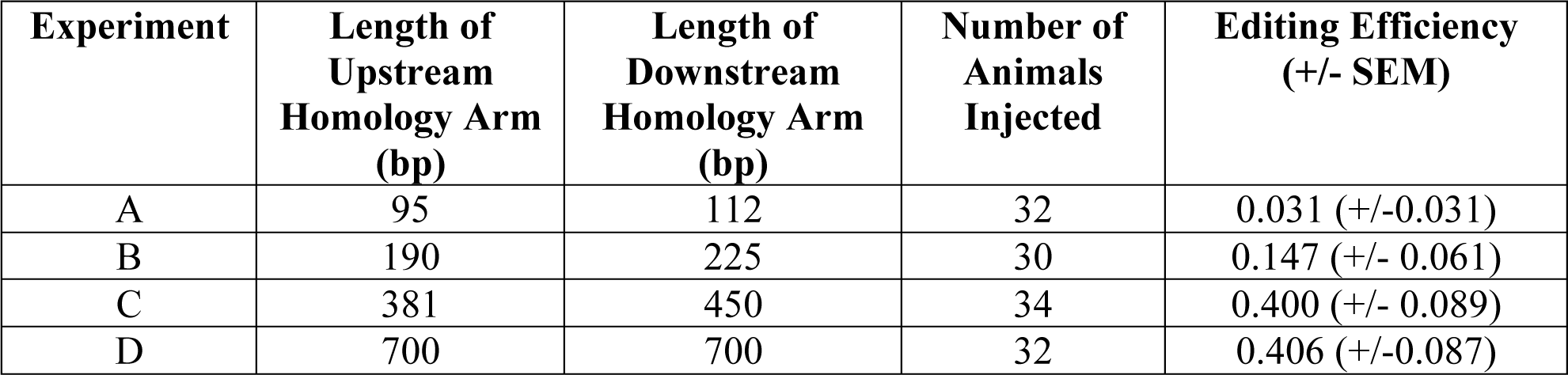
Editing efficiency for a range of tested homology arm lengths.

| **Experiment** | **Length of Upstream Homology Arm (bp)** | **Length of Downstream Homology Arm (bp)** | **Number of Animals Injected** | **Editing Efficiency (+/- SEM)** |
| --- | --- | --- | --- | --- |
| A | 95 | 112 | 32 | 0.031 (+/-0.031) |
| B | 190 | 225 | 30 | 0.147 (+/- 0.061) |
| C | 381 | 450 | 34 | 0.400 (+/- 0.089) |
| D | 700 | 700 | 32 | 0.406 (+/-0.087) |

For this experiment, we followed the CRISPR protocol described in Materials and Methods with one modification: injected P0s were singled onto plates in order to obtain an accurate measure of editing efficiency, which was defined as the proportion of P0 plates bearing progeny with the selection cassette integrated at the desired location, as confirmed by PCR. The highest editing efficiency was obtained when utilizing homology arms over approximately 400 bp (see experiments C and D, Figure 3). When homology arms smaller than these were used, as in experiments A and B, the editing efficiency dropped significantly (Figure 3). These results agree with previous findings (Dickinson *et al*. 2015; Norris *et al*. 2015), suggesting that the optimal length of homology for efficient genome editing is dependent on the particular application of HDR being used. As we observe no further gain in efficiency of HDR for larger deletions with homology arms after about 400 bp, we have adjusted our protocol accordingly. For our current production protocol, we order homology arms as 500-bp gBlocks from IDT with 50 bp of adaptor sequences included to enable Gibson assembly (Gibson *et al*. 2009) into our repair vector.

**Figure 3.**
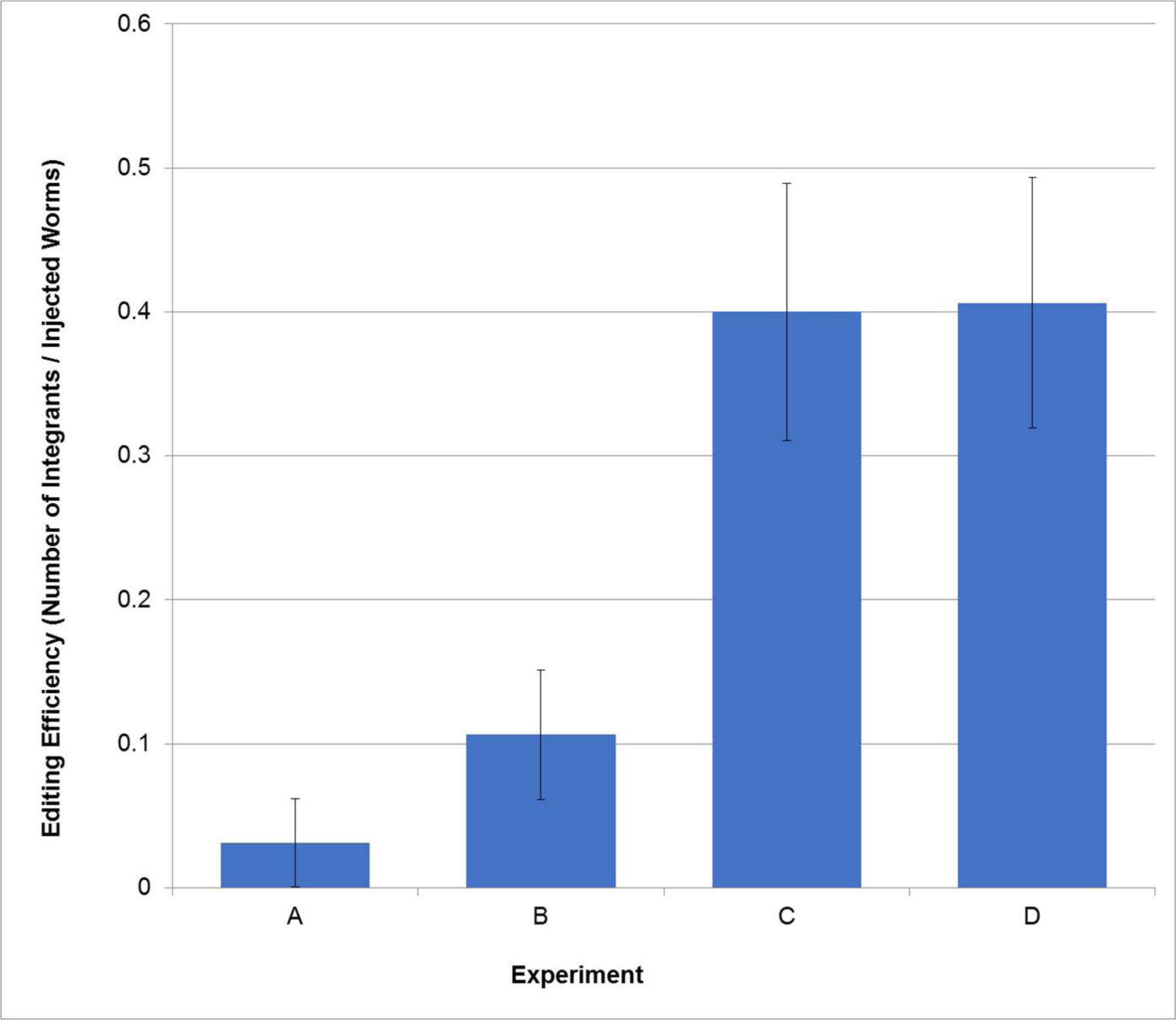
Assessing editing efficiency with homology arms of varying lengths. A 536-bp deletion was generated using a single guide RNA and various lengths of homology arms (see Table 1 for details). The proportion of individual injected P0s giving rise to animals with the selection cassette integrated at the desired location in the genome was determined by PCR validation. Error bars indicate standard error of the mean.

### Delivery of purified Cas9 protein versus plasmid-borne Cas9

There are several reports in the literature from researchers using mammalian systems indicating that efficiency of Cas9 cutting improves if they use purified Cas9 protein rather than plasmid-borne Cas9 (reviewed in Wu *et al*. 2014). Results confirming these observations have been reported for *C. elegans* (Paix et al. 2014). There is some debate on how much Cas9 protein is required for making a DSB in this organism. All results reported here were achieved using Cas9 protein at a concentration between 1.5 µM to 3 µM. Our results confirm that, as in other systems and as previously reported for this nematode, purified Cas9 is more efficient at inducing DSBs than plasmid-borne Cas9. Our conclusions are based on testing 112 gene targets (Figure 4A). In cases where we have tested the same gene with both purified protein and plasmid-borne Cas9 (three genes), using the purified protein is four times more efficient at inducing a double-strand break at the target site (Figure 4B). There are also reports from other model systems that conclude the use of purified protein led to fewer non-specific effects than the plasmid. As discussed below, that is not the case for *C. elegans*, where the two delivery methods exhibit an identical low level of non-specific effects.

**Figure 4.**
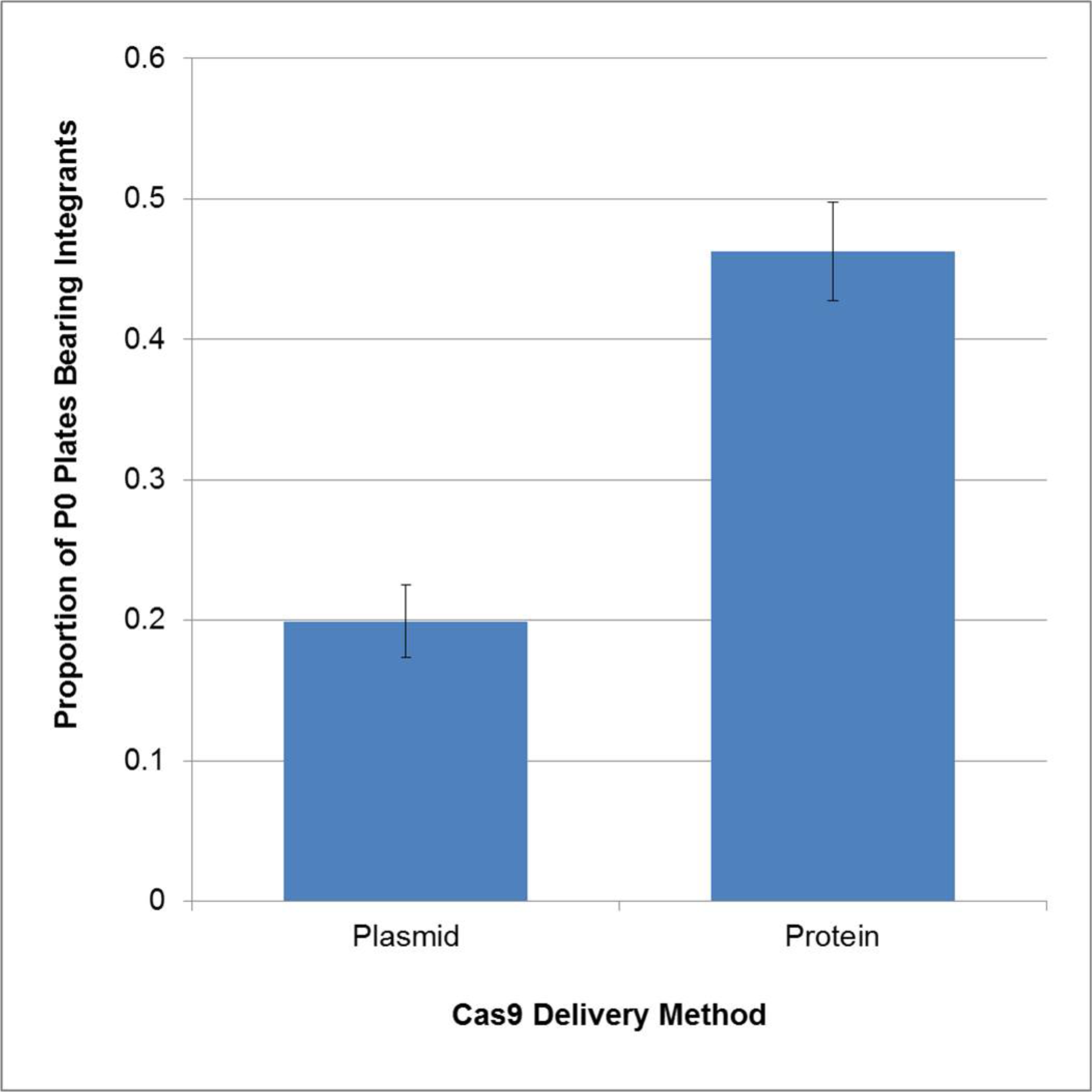
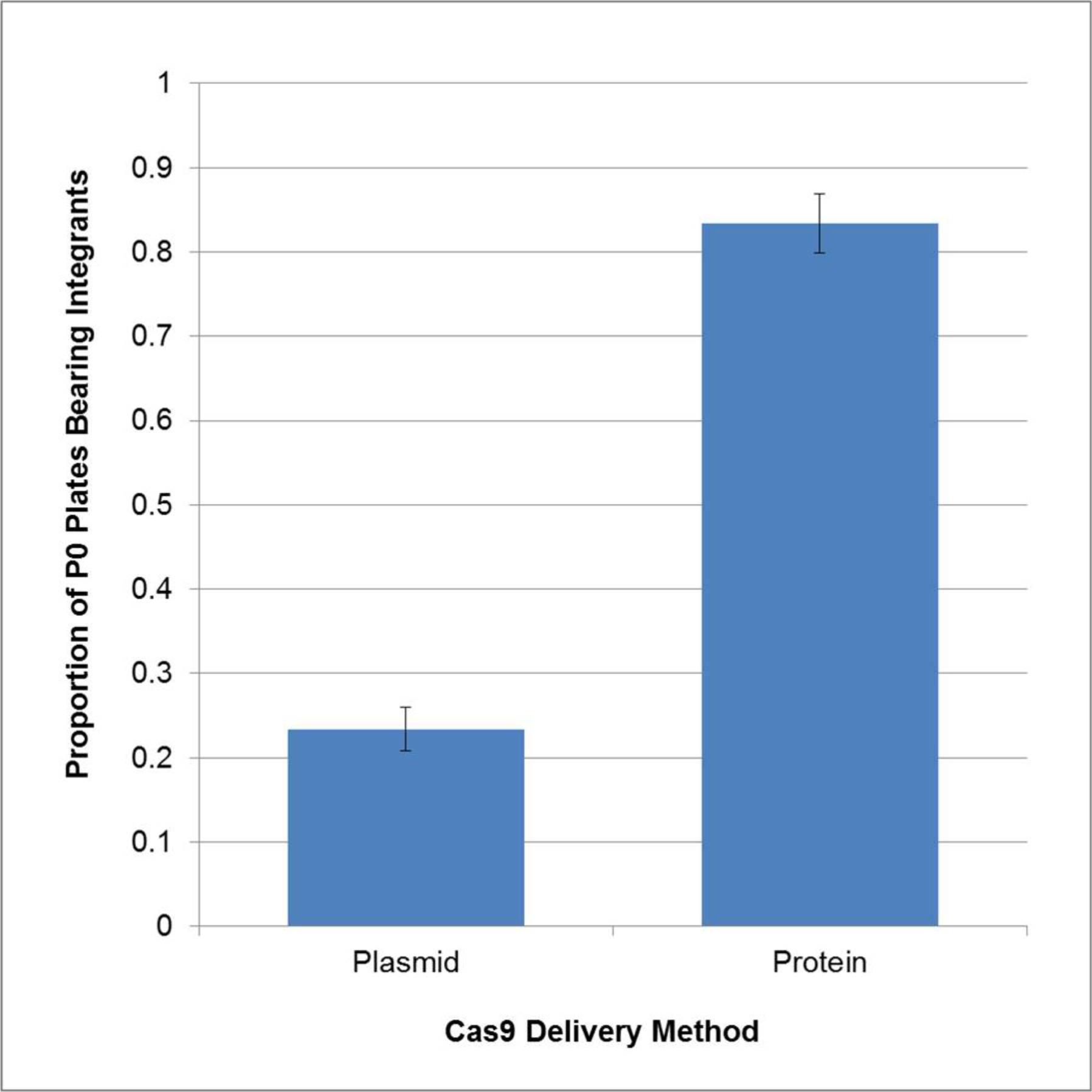
A and B. Comparison of Cas9 delivery methods. For 112 genes targeted with different delivery methods of Cas9, the proportion of P0 plates (4 injected animals per plate) bearing progeny with the selection cassette integrated at the desired location was determined by PCR validation. (A) Animals injected with the plasmid-borne Cas9 (N=56) were less efficient at producing integrants than those injected with purified Cas9 protein (N=59). (B) A direct comparison of editing efficiency in which the same three gene targets were tested both with Cas9 plasmid and protein. Error bars indicate standard error of the mean.

### Off-target effects

At present there appear to be conflicting results in the literature with regard to off–target events when using the CRISPR/Cas9 gene editing system. Whether such events are due to guide RNA homology, homology of the repair template, or unrelated spurious Cas9 cutting is unclear. As it is critical that we know of any such events in the strains we provide to the CGC and ultimately to the larger worm community we undertook a series of tests to determine the frequency of off-target events after CRISPR/Cas9 treatment.

In our study we performed WGS on eight CRISPR/Cas9-derived strains and their parental strain (see Table 2). For this experiment we targeted two distinct genes using two different Cas9 delivery methods. We searched for small off-target mutations, both single nucleotide variants (SNVs) and indels, using strict filtering in order to eliminate variants already present in the parental population as well as potential technical artefacts (see Methods). We found a total of eight mutations (see Table 2), six homozygous and two heterozygous calls. None of the mutations appear to be related to either the guide RNAs or the homology arms. We also searched for off-target genomic rearrangements by analyzing copy number estimates (see Methods and Itani *et al*, 2015), but none were found. The frequency of SNVs and indels detected is on the order of spontaneous mutations reported for this organism (Denver *et al*. 2009), which suggests the observed mutations are unlikely to be due to the CRISPR/Cas9 procedure. In our quality control studies we have used WGS to analyze an additional 30+ CRISPR/Cas9-generated deletion lines, and again, we found little or no evidence for off-target events (data not shown). We conclude that off-target mutations due to this method are rare and do not appear to be a serious concern in *C. elegans* when guide RNAs are carefully selected. We also conclude that in this nematode, unlike what has been reported for other systems, the delivery method (i.e. plasmid *versus* protein) does not appear to influence the occurrence of off-target mutations.

**Table 2.** Whole-genome sequencing statistics for the parental strain and 8 mutant strains produced for the off-target mutation analysis.

| **Strain** | **Target Gene** | **Cas9 Delivery** | **Read Length (bp)** | **Mean Coverage** | **Number of off-target SNVs** | **Number of off-target indels** |
| --- | --- | --- | --- | --- | --- | --- |
| VC3823 | lgc-45 | plasmid | 1x75 | 22 | 2 | 0 |
| VC3821 | lgc-45 | plasmid | 2x300 | 34 | 0 | 0 |
| VC3817 | lgc-45 | protein | 2x75 | 34 | 0 | 0 |
| VC3814 | lgc-45 | protein | 2x75 | 30 | 0 | 0 |
| VC3843 | C34D4.2 | plasmid | 2x75 | 27 | 1 | 0 |
| VC3845 | C34D4.2 | plasmid | 2x75 | 34 | 1 | 0 |
| VC3834 | C34D4.2 | protein | 2x75 | 32 | 2 | 1 |
| VC3840 | C34D4.2 | protein | 2x75 | 30 | 1 | 0 |
| VC3504 | parent | - | 2x300 | 41 | - | - |

### Validating deletion events at the target site

While we detected virtually no off-target events, we now have several examples where target deletions are accompanied by local rearrangements, including local duplications, deletions of adjacent DNA, or insertions of multiple copies of the repair template (see for example Figure 5). This is data we accumulated after WGS of more than 30 CRISPR/Cas9 derived strains for several target genes. Perhaps it should be of no surprise that a double-strand break, while it can yield HDR, could also be resolved in other ways. As this will be a recurring problem and does not seem to be specific to the gene, guide RNA, or homology arms, we have developed a Quality Control (QC) deletion validation protocol (see Figure 6 and Table 3). Our initial selection is based on drug resistance and Mendelian segregation of weak GFP expression in the pharynx. This is strong evidence that we have targeted and disrupted the gene of interest, but it does not guarantee that we have not disrupted flanking DNA, or even indicate whether we still have an intact copy of the target gene. Our PCR-based QC protocol tests for these possibilities by verifying the integrity of the disruption junction on both sides of the insert and confirming that the wild type sequence is indeed absent. Primer pairs chosen and expected results are shown in Table 3. Our experience to date tells us that we should expect only a portion of drug-selected GFP positive animals to be correctly edited for any particular gene and for this reason we try to obtain at least 5 or 6 positive integrants per target gene.

**Figure 5.**
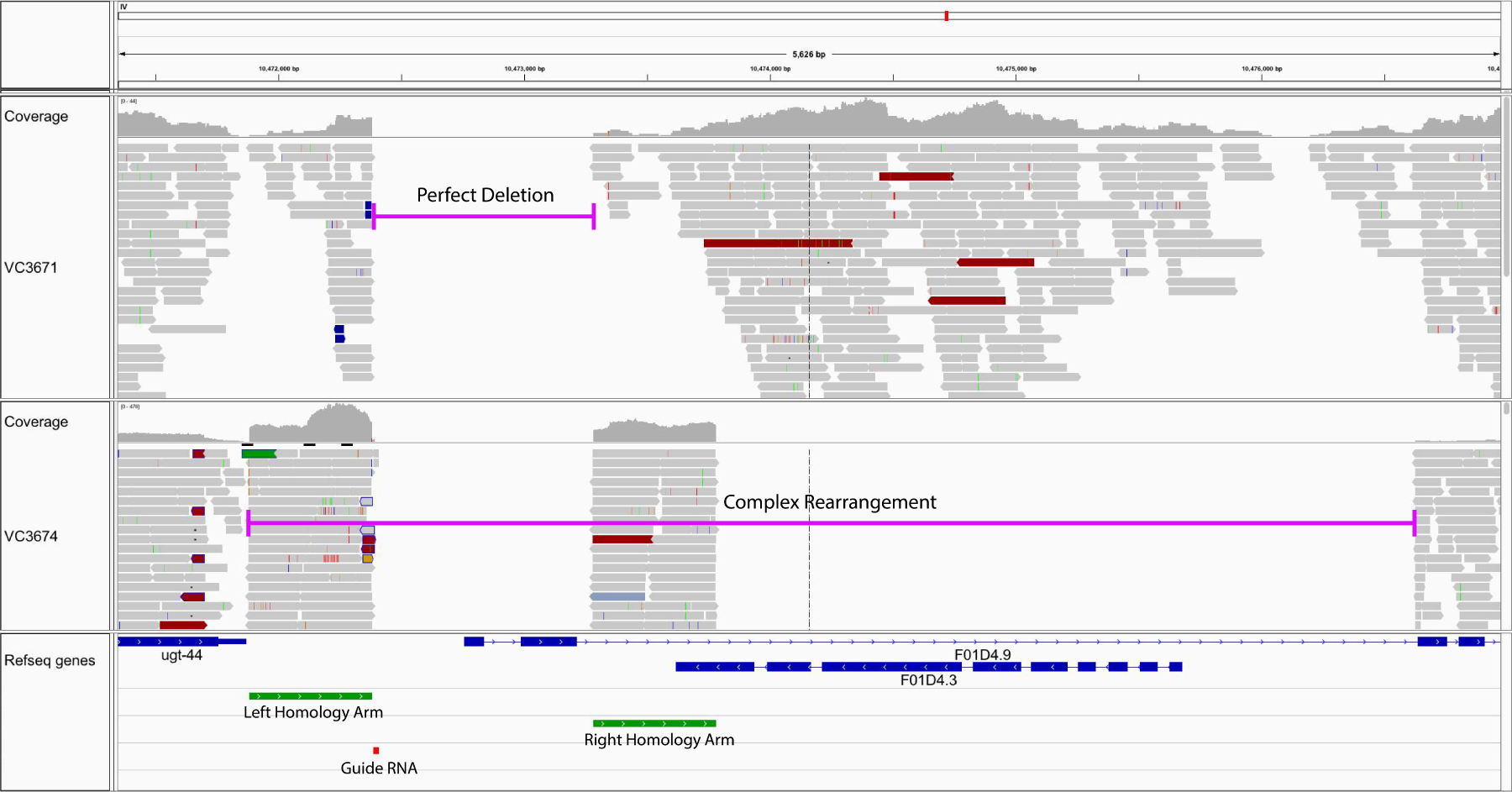
Rearrangements at the target site. Two examples of removing the target gene through CRISPR/Cas9 HDR aligned using IGV. In strain VC3671 there is a perfect deletion removing the first two exons of gene F01D4.9. In strain VC3674 there is evidence of a complex rearrangement. There is a deletion of the same two exons as well as the downstream region of F01D4.5 and a pseudogene, F01D4.3. The deletion extends much further downstream and likely harbours multiple copies of the repair template, since the average coverage in the homology arm regions is relatively high compared to the adjacent genomic sequence. In blue are the exons for the various genes. In green are the homology arms chosen for F01D4.9 and in red is the guide RNA.

**Figure 6.**
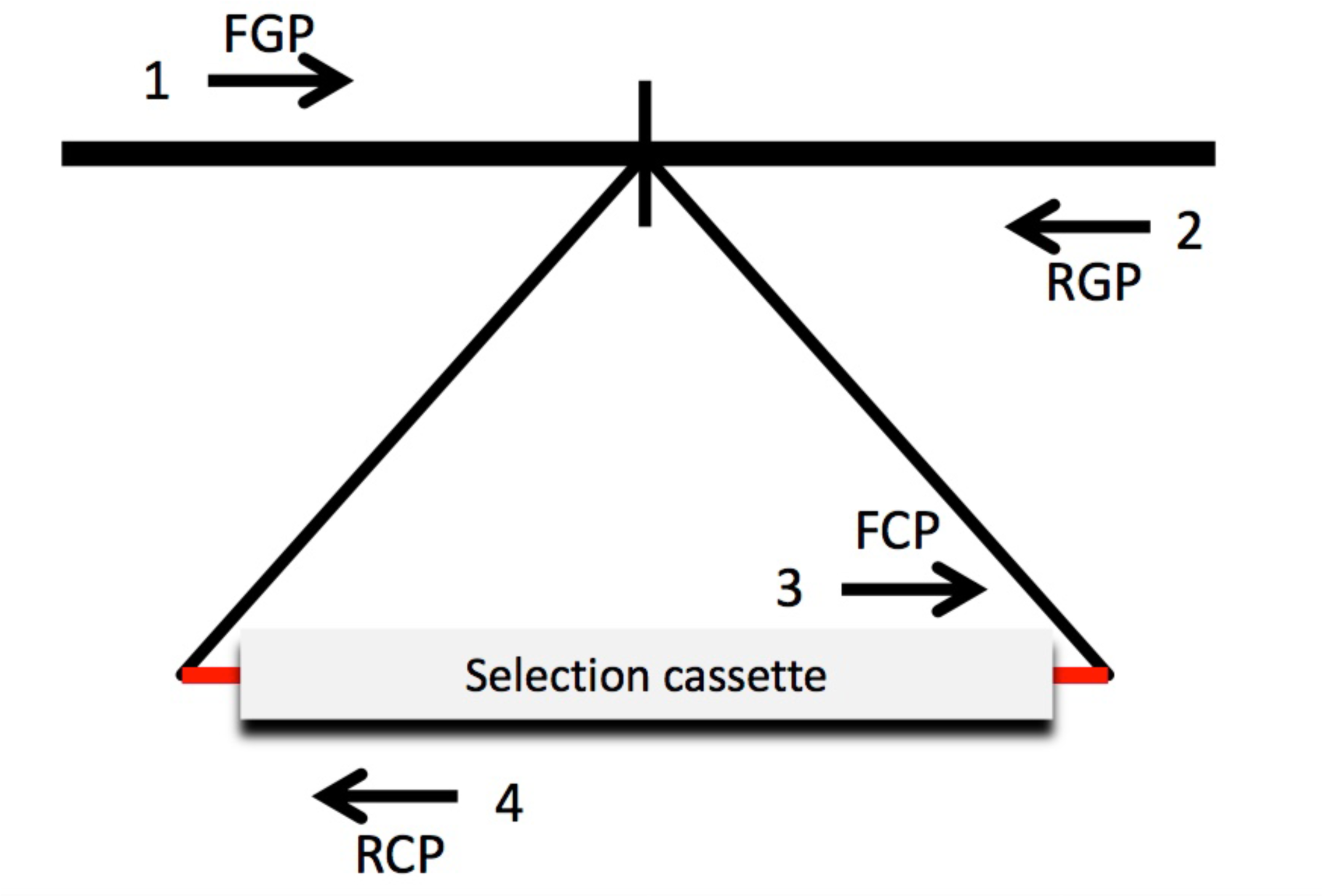
Graphical representation of Quality Control (QC) Validation Scheme. A typical CRISPR/Cas9-generated edit is represented by a deletion (dotted line) of genomic sequence being replaced by the selection cassette below. Arrows indicate the position of primers for QC PCR validation (see Table 3). FGP: forward genomic primer. RGP: reverse genomic primer. FCP: forward cassette primer. RCP: reverse cassette primer.

**Table 3.** Quality Control (QC) PCR Interpretation.

| Diagnostic State | Left Insertion PCR (Primers 1+2) | Right Insertion PCR (Primers 3+4) | WT PCR on Mutant (Primers 1+4) | WT PCR on WT (Primers 1+4) | Conclusion |
| --- | --- | --- | --- | --- | --- |
| A | Correct | Correct | No product | Correct | Mutant, probably as expected |
| B' | Correct | Incorrect | No product | Correct | Mutant with rearrangement |
| B'' | Incorrect | Correct | No product | Correct | Mutant with rearrangement |
| C | Incorrect | Incorrect | No product | Correct | Mutant with rearrangement |
| D | Incorrect | Incorrect | WT product | Correct | Not mutant |

## Discussion

We have investigated several variables that may impinge on the effectiveness of the CRISPR/Cas9 procedure to generate large deletions in *C. elegans*. Our *C. elegans*- specific guide RNA selection tool in conjunction with the selection vector designed by the Calarco group (Norris *et al*. 2015), and the use of purified Cas9 protein, as pioneered by the Seydoux group for use in this organism (Paix *et al*. 2016), yields a very efficient and effective protocol. We feel this protocol has sufficient robustness to tackle the remaining genes in this organism that lack null alleles.

Among the parameters we explored in refining an HDR protocol appropriate for our purposes were homology arm length and Cas9 delivery methods, i.e. delivery via plasmid or as a purified protein in a nucleoprotein complex. These parameters were previously explored (Dickinson *et al*. 2015; Norris *et al*. 2015; Paix *et al*. 2014, 2016), but in somewhat different conditions than those reported in this study. The length of the homology arm required is at least partially dependent on the type of edit being done to the gene. For single nucleotide changes or small indels, as little as 35 nucleotides of homology are required when using single-stranded bridging oligonucleotides (ssODNs) (Paix *et al*. 2016). As our facility is looking to make larger deletions that remove most or all of an open reading frame (ORF) this approach is not feasible. For this reason, we were more attracted to protocols developed for replacing larger stretches of DNA. We also preferred protocols that replaced screening DNA directly and instead use a drug and/or visible marker for selection as we envision performing hundreds of screens using the protocol. As stated earlier, we examined the Dickinson *et al*. (2015) and Norris *et al*. (2015) vectors and settled on using the G418-resistant, pharyngeal GFP selection system of Norris *et al*. (2015). We chose the latter system because it has a smaller selection cassette. Both of these groups agree that longer homology arms are required for making large insertions/deletions via HDR. They both experimented with homology arms of one to two kb but also pointed out that 500 bp arms would work. We concur with these findings and extend these studies to show there is no advantage to making arms longer than 500 bp. With homology length of only 450 bp, we were able to generate up to 20kb deletions (using two guide RNAs – data not shown). From a recent paper using CRISPR/Cas9 to generate inversions in *C. elegans* it appears there is no limit to the spacing of simultaneous cut sites (Dejima *et al*. 2018).

Similar to Paix *et al*. (2016), we find that purified protein is more efficient at generating on-target DSBs. In our hands the purified protein is at least four times more effective than plasmid-borne Cas9 when tested against numerous genes. There are a number of possible reasons for this difference but the most likely one is that there is substantial lag between injecting a plasmid and producing a protein, which then has to be combined with the guide RNA. This lag time is removed when injecting an RNP complex directly. Overall protein concentration between the two methods may also differ.

We detect virtually no off-target events after using our refined CRISPR/Cas9 protocol. We suspect the major contributor to eliminating off-target events is the use of our *C. elegans*-specific guide RNA selection tool. Several reviews on the issue of off-target effects consider poor guide RNA design to be a major contributor to these effects (see for example Wu *et al*. 2014 and Jamal *et al*. 2015). Additionally, while we saw no difference in the frequency of off-target events between Cas9 delivery systems, others have shown that in mammalian cells, direct delivery of Cas9 protein and guide RNA as a nucleoprotein complex reduces off-target events (Kim *et al*. 2014; Ramakrishna *et al*. 2014). Considering that the deletion lines we generate will go to the CGC for distribution to the nematode community, identifying and eliminating the sources of off-target events is imperative. It seems the protocol we have developed satisfies this criterion.

## Acknowledgements

Dr. John Calarco (University of Toronto) and Dr. Geraldine Seydoux (Johns Hopkins University) kindly provided us with reagents and much valued advice as we refined the CRISPR/Cas9 system in our laboratory. We also thank Mei Zhen and Julie Claycomb, for their critical support during the early stages of these studies. This work was funded by CIHR grant PJT-148549 to DGM, CIHR grant MOP 93719 and NSERC grant 312498 to HH, NIH grant 5P40OD010440 to AR and NIH 5R240D023041 grant to AR, Paul Sternberg, Geraldine Seydoux and DGM.

